# Cerebrovascular health mediates processing speed change through anterior white matter alterations: A UK Biobank Study

**DOI:** 10.1101/2024.05.10.593527

**Authors:** KL Moran, CJ Smith, E McManus, SM Allan, D Montaldi, N Muhlert

## Abstract

Cerebrovascular disease is associated with an increased likelihood of developing dementia. While cardiovascular risk factors are modifiable and may reduce the risk of later-life cognitive dysfunction, the relationship between cerebrovascular risk factors, brain integrity and cognition remains poorly characterised. Using a large UK Biobank sample of predominantly middle-aged adults, without neurological disease, our structural equation mediation models showed that poor cerebrovascular health, indicated by the presence of cerebrovascular risk factors, was associated with slowed processing speed. This effect was best explained by anterior white matter microstructure changes, rather than posterior changes. Effects were also significantly reduced when considering other forms of cognition, demonstrating both regional- and cognitive-specificity of our effects. Critically, our findings also demonstrate that including measures of risk factor duration may be particularly important for improving estimations of cerebrovascular burden. In summary, our study demonstrates the specific impact of early cerebrovascular burden on brain structure and cognitive function, highlighting the necessary next steps for improving cerebrovascular burden quantification and improving clinical predictions.

## Introduction

Cerebrovascular disease is closely linked with the development of later-life cognitive dysfunction and dementia ^1 2^. Many of the risk factors associated with cerebrovascular disease are modifiable, such as hypertension and smoking status, which suggests they may be an optimal group of therapeutic targets with which to minimise or even prevent later cognitive dysfunction. However, in order to identify the risk factors which will yield maximal impact when targeted, a mechanistic understanding of the relationship between cerebrovascular health and cognition is needed; particularly during the earlier stages of this relationship, where intervention is more likely to have a beneficial impact on both brain integrity and cognitive function.

White matter is particularly vulnerable to vascular insult, due to its cellular composition and absence of collateral circulation^3^ and is likely key to understanding how cerebrovascular burden influences cognition. Macroscale markers of white matter injury, such as white matter hyperintensities (WMHs) are consistently linked to both poor vascular health and cognitive dysfunction^4^. However, WMHs reflect an advanced stage of white matter injury and it’s unlikely that intervention at this stage would have a beneficial impact on brain health or cognitive function^5^. Diffusion imaging allows for microscale white matter changes to be mapped, which reflect a much earlier stage of injury that may be more receptive to intervention, relative to their macroscale counterparts^6^. These microscale measures of white matter integrity (WMI) may be particularly valuable when applied to studying middle-age populations in the early stages of cerebrovascular burden, where damage is likely subtle and difficult to detect. WMI is shown to correlate well with cerebrovascular health and individual risk factors, such as hypertension ^7^, diabetes ^8 9^ and obesity ^10^, and is sensitive in predicting stroke risk ^11^. Therefore, it’s likely that subtle WMI changes may act as a sensitive biomarker of early cerebrovascular burden.

A growing body of evidence suggests the impact of cerebrovascular burden on white matter may be region-specific. More specifically, the differences in the vascular characteristics of anterior and posterior white matter regions may moderate the impact of cerebrovascular burden on the brain, particularly in its early stages. For example, hypertension is associated with greater change in anterior cerebrovascular function, such as reduced cerebral blood flow, relative to posterior regions ^12 13^. This suggests a unique vulnerability of anterior regions to impaired vascular health. Region-specific white matter damage may also explain some of the inconsistent findings in the literature. For example, one study found no correlation between superior longitudinal fasciculus (SLF) WMI and cerebrovascular burden ^14^. However, using the same UK Biobank sample, albeit with small differences in exclusion criteria, another study found a strong relationship between WMI and cerebrovascular burden ^15^. The latter study also evidenced some anterior-specific effects of cerebrovascular burden, which were likely not sufficiently captured by the SLF alone in the former study. In addition, the former study used an aggregate method to quantify cerebrovascular burden, which may inflate the effects of noise and underestimated effects ^14 16^. A large sample study investigating regionally specific effects of cerebrovascular burden, which adopts sufficiently sensitive metrics and quantification methods is necessary to address these shortfalls.

Given the proposed regional-specific impact of cerebrovascular burden, it follows that there may be some specificity to the cognitive consequences of this burden. Cerebrovascular disease has been linked with deficits in the cognitive processes supported by frontal regions, including processing speed, executive function and working memory ^17 18 19 20^. However, processing speed is most consistently linked to cerebrovascular health and anterior white matter change ^21^. Salthouse proposed that slowed processing speed precedes decline in other cognitive domains ^22^. While it remains unclear whether this is the case with cerebrovascular burden in a middle aged cohort, evidence indicates that processing speed mediates the impact of vascular burden on other cognitive domains ^23^. Processing speed may therefore be particularly well suited as a candidate early marker of cerebrovascular burden. This is further supported by evidence of a strong link between WMI and slowed processing speed ^21^. We can therefore postulate a mechanistic pathway through which cerebrovascular burden alters white matter, and manifests behaviourally as subtle, but accumulating, cognitive change, likely beginning with alterations in speed of processing.

The *duration* of cerebrovascular risk factors may also significantly impact upon brain integrity and cognition. Studies reporting links between cerebrovascular burden, cognition and brain health traditionally quantify cerebrovascular burden based on the summed presence of known risk factors at the time of assessment ^14 15^. However, this static approach may underestimate cerebrovascular burden, and consequently, its impact. Using this approach, an individual with a chronic history of hypertension would be considered to yield the same degree of burden as an individual with a recent diagnosis. This is inherently problematic, as the duration of hypertension is significantly exacerbates both white matter alterations ^24^ and cognitive dysfunction ^25^, and in particular, processing speed ^26^. Additionally, smoking and diabetes duration are also thought to contribute to white matter change over time, but the cognitive impact of risk factor duration is not fully understood ^27 9^.

While the evidence to date suggests a viable mechanistic pathway through which early cerebrovascular burden may lead to cognitive impairment, further research is needed to characterise and better understand the specificity of this pathway. This will improve our understanding of cerebrovascular cognitive health from a systems perspective, and inform the development and use of appropriate and sensitive cognitive assessments for the early detection and monitoring of cerebrovascular cognitive decline. With this ambition in mind, the aims of the present study were twofold. First, we aimed to explore the relationship between cerebrovascular burden, WMI and cognition in middle-aged participants and investigate whether any effects of cerebrovascular burden were specific to anterior WMI or to processing speed. We predicted that as cerebrovascular burden increases, processing speed will slow and that this effect will be mediated by anterior WMI. Furthermore, we also predicted that these effects would be specific to, or at least more pronounced for, processing speed, compared to other domains of cognition and to anterior WMI, rather than posterior WMI. Our second aim was to assess whether processing speed was better predicted by cerebrovascular burden when accounting for risk factor duration, as opposed to models of static cerebrovascular burden.

## Results

### Demographic Information, Descriptive Statistics & Factor Loadings

Cerebrovascular, neuroimaging and cognitive data were analysed in a sample of 37,265, aged 40-70, derived from the UK Biobank. Descriptive statistics for demographic information and cerebrovascular risk factors are described in *Table 1*. Data shown are described prior to being transformed (e.g. BMI - log transformed) and scaled (all data). All variables loaded significantly (*p <* .*001)* onto their respective latent construct, which are depicted in *Figure 1 and Tables 2 & 3, respectively*.

**Table 1.** Demographic & cerebrovascular risk factor descriptive statistics. Continuous variables are represented as mean (standard deviation) and binary variables are given as a percentage.

| Variable | Unit | Descriptor |
| --- | --- | --- |
| **Demographics** |  |  |
| Age | Years, M(SD) | 65.41 (7.66) |
| Sex | Female (%) | 53.30% |
| Ethnicity | White British (%) | 91.04% |
| **Risk factors** |  |  |
| Smoking (current or past) | Yes (%) | 36.79% |
| Diabetes | Yes (%) | 5.05% |
| High cholesterol medication | Yes (%) | 22.14% |
| BMI | Kg / m<sup>2</sup>, M(SD) | 26.45 (4.35) |
| Systolic (BP) | mmHG, M(SD) | 138.64 (18.61) |
| Diastolic (BP) | mmHG, M(SD) | 78.72 (10.02) |
| Waist to Hip Ratio | W:H, M(SD) | 0.88 (0.09) |
| Smoking duration | Years, M(SD) | 20.07 (16.62) |
| Diabetes duration | Years, M(SD) | 11.62 (10.06) |
| Hypertension duration | Years, M(SD) | 13.72 (8.61) |

**Table 2.** Factor loadings for anterior white matter integrity latent construct. Table contains loadings for anterior tracts for both fractional anisotropy (FA) and mean diffusivity (MD).

| Anterior White Matter Tract | Hemisphere | Factor loading |  |
| --- | --- | --- | --- |
|  |  | FA | MD |
| Genu of Corpus Callosum | - | .759* | .775* |
| Anterior limb of Internal Capsule | Left | .682* | .787* |
|  | Right | .665* | .754* |
| Anterior Corona Radiata | Left | .809* | .911* |
|  | Right | .771* | .922* |
| Superior Corona Radiata | Left | .662* | .807* |
|  | Right | .639* | .790* |
| Superior Longitudinal Fasciculus | Left | .702* | .853* |
|  | Right | .715* | .839* |
| Superior Fronto-occipital Fasciculus | Left | .568* | .629* |
|  | Right | .541* | .671* |
| Uncinate Fasciculus | Left | .513* | .430* |
|  | Right | .488* | .503* |

Note: \* $p < .001$

**Table 3.** Factor loadings for posterior white matter integrity latent construct. Table contains loadings for posterior tracts for both fractional anisotropy (FA) and mean diffusivity (MD).

| Posterior White Matter Tract | Hemisphere | Factor loading |  |
| --- | --- | --- | --- |
|  |  | FA | MD |
| Splenium of Corpus Callosum | - | .667* | .715* |
| Pontine Crossing Tract | - | .266* | .178* |
| Middle Cerebellar Peduncle | - | .625* | .463* |
| Cerebral Peduncle | Left | .683* | .439* |
|  | Right | .689* | .468* |
| Inferior Cerebellar Peduncle | Left | .602* | .481* |
|  | Right | .631* | .512* |
| Superior Cerebellar Peduncle | Left | .563* | .476* |
|  | Right | .560* | .473* |
| Posterior Corona Radiata | Left | .747* | .814* |
|  | Right | .714* | .803* |
| Sagittal Striatum | Left | .685* | .804* |
|  | Right | .710* | .827* |

Note: \* $p < .001$

**Figure 1.**
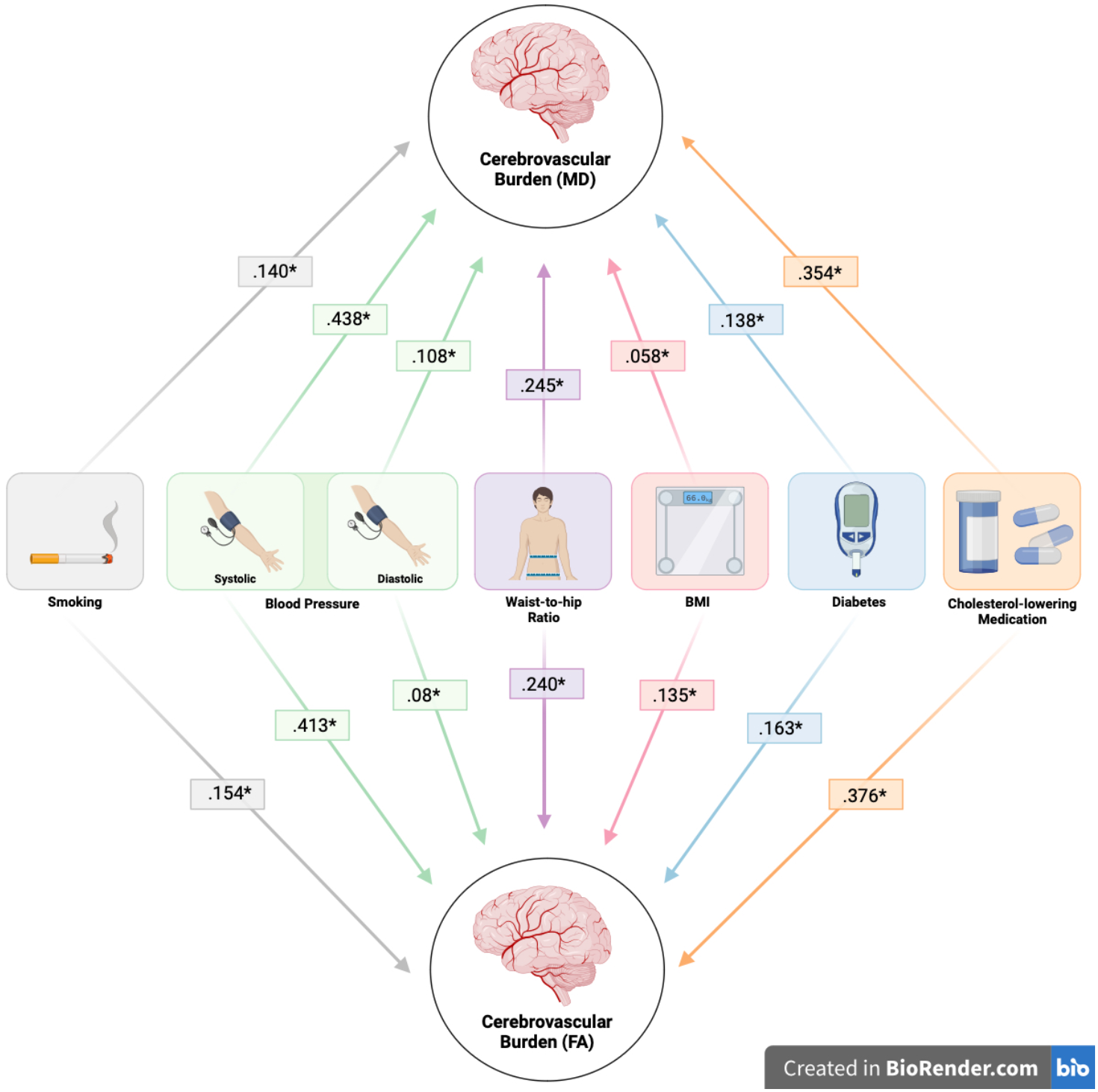
Factor loadings for each cerebrovascular risk factor onto the latent construct of ‘Cerebrovascular Burden’ in separate Fractional Anisotropy (FA) and Mean Diffusivity (MD) models. Each risk factor loaded significantly (*p < .001). This figure was created using BioRender.

### Principal Component Analyses (PCA)

A PCA derived two principal components, which were subsequently used as latent constructs in the SEMs. The two components and their variable loadings are illustrated by the variable correlation plot in *Figure 2*. Here, positively correlated variables are grouped together, with the *X* and *Y* axis representing principal component 1 and 2, respectively. The quality of representation of the variables on the factor map is represented by the colour gradient (squared cosine coordinates – ‘*cos2’*), this illustrates how well a variable reflects a given component. Variables that are coloured in a darker shade of the gradient spectrum reflect a high value, indicating that the variable is a good representation of the component.

**Figure 2.**
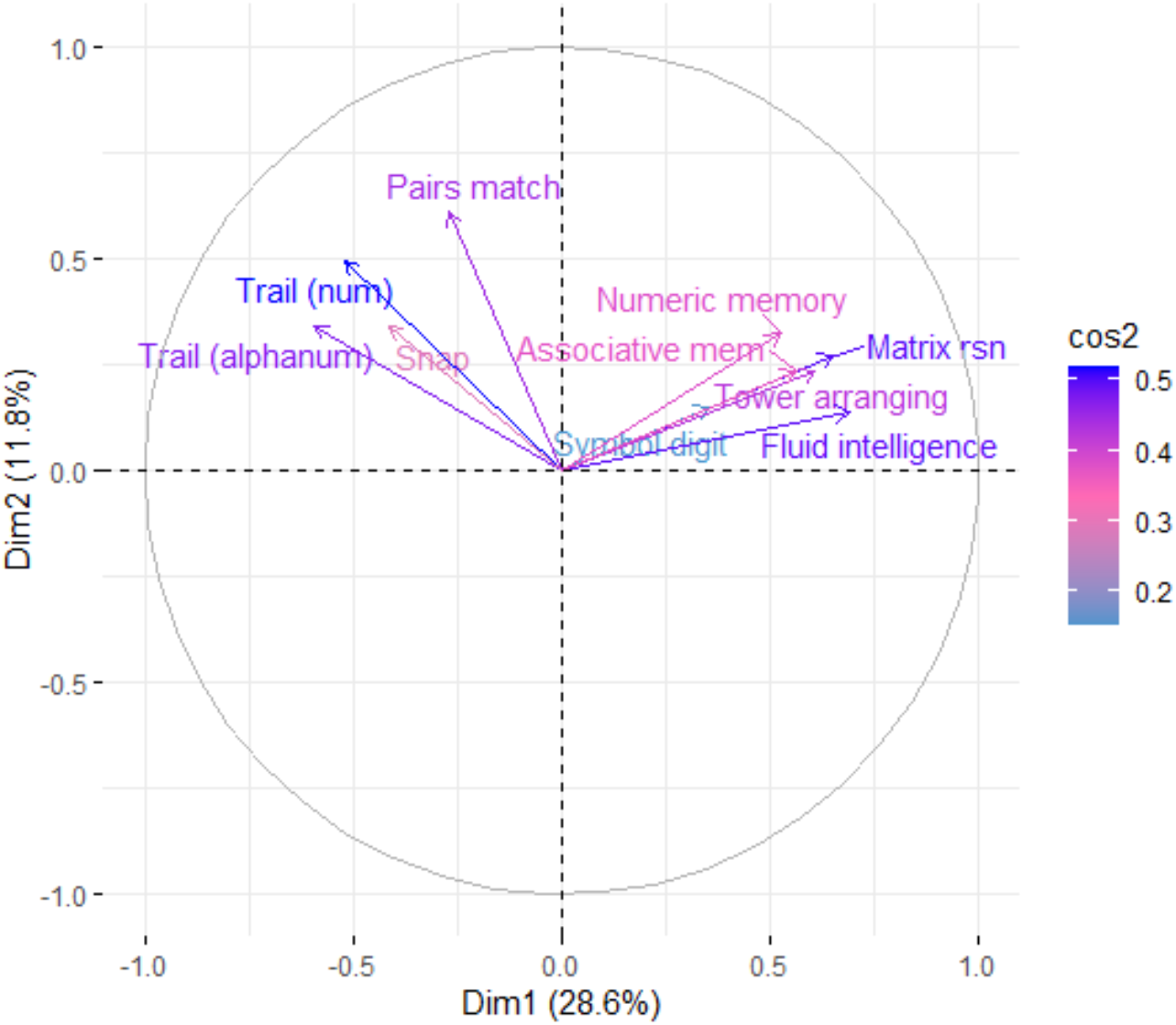
Correlational plot illustrating the relationship between individual cognitive tasks and the first 2 principal components and the quality of variable representation. Principal component 1 is represented on the X-axis and principal component 2 is represented by the y-axis. The gradient colour bar reflects the squared cosine coordinates, which indicates the quality of representation of each cognitive task on their respective principal component. Figure was created using ‘FactoExtra’ package in R.

The contribution of each task to principal components 1 & 2 is shown in *Figure 3*. Based on this analysis and the pattern of task-types associated with each principal component, we suggest principal component 1 reflects general cognitive function, while principal component 2 reflects processing speed. We drew this conclusion by examining the tasks and measures most highly associated with each principal component. For example, the tasks most highly associated with principal component 1 are Fluid Intelligence, Matrix Reasoning and Tower Arranging. These tasks reflect general cognitive function, as they draw on a number of different cognitive processes, such as abstract reasoning, attention and spatial perception and these are reflected by performance metrics. In comparison, the tasks that loaded most highly on principal component 2 are Pairs, Trail Making (numeric) and Snap. For each of these tasks, processing speed is the dominant cognitive process, and speed metrics were derived from them as measures of performance.

**Figure 3.**
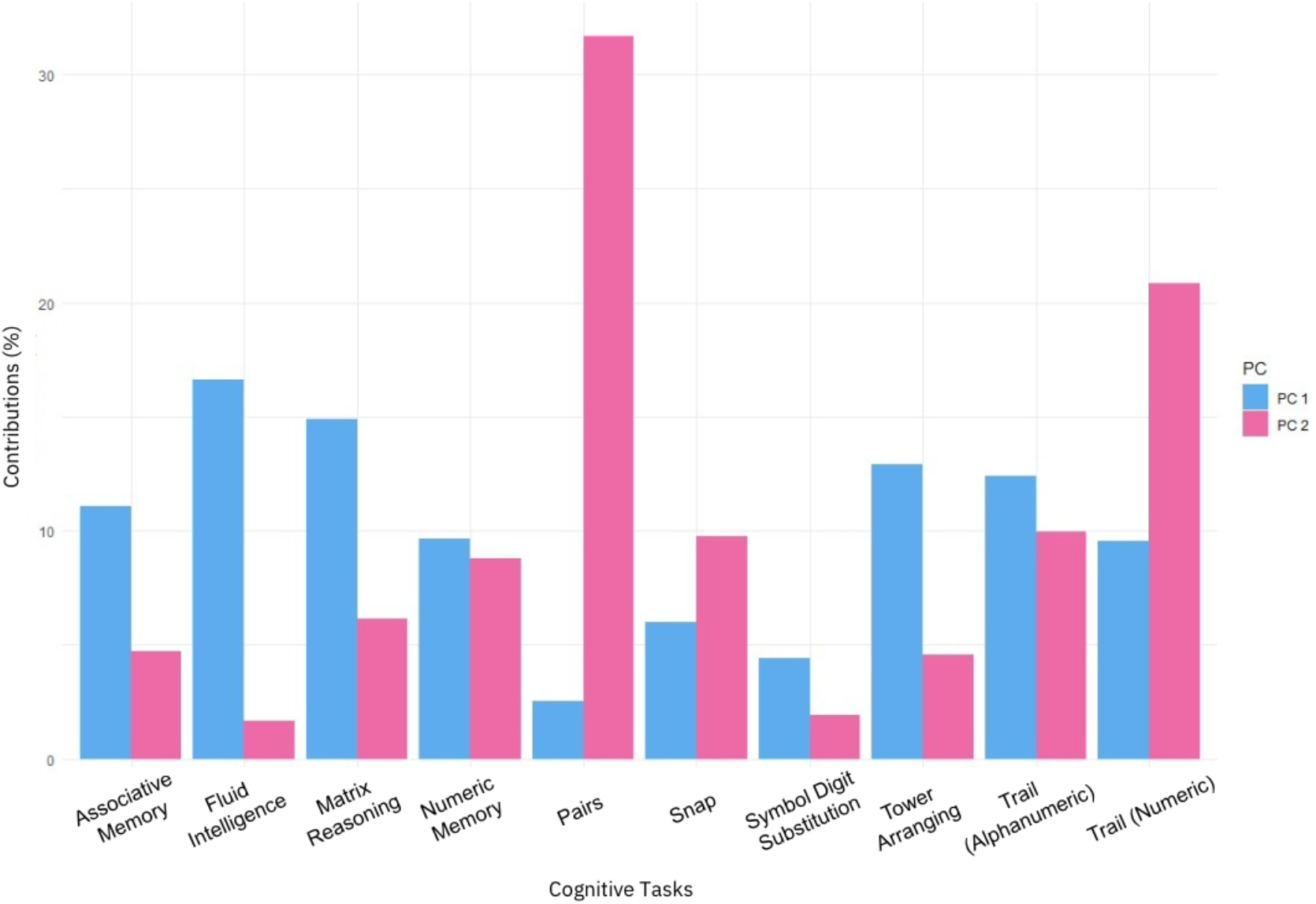
Contributions (%) of each individual cognitive task to principal component 1 & 2. Principal component 1 & are depicted in blue and pink, respectively.

### Structural Equation Modelling (SEM)

Once both the latent construct for cerebrovascular burden and anterior WMI were created, they were loaded into a mediation model alongside the cognitive processing speed principal component. Here, a significant direct effect was found, with increased cerebrovascular burden associated with slowed cognitive processing speed (*p* < .001). The indirect effect of this model was also significant, demonstrating that anterior WMI loss partially mediated the relationship between cerebrovascular burden and processing speed (*p* < .001). Model fit indices indicating goodness of fit are shown in *Figure 4a*, alongside path and significance values.

**Figures 4. a (left), 4b (middle) & 4c (right).**
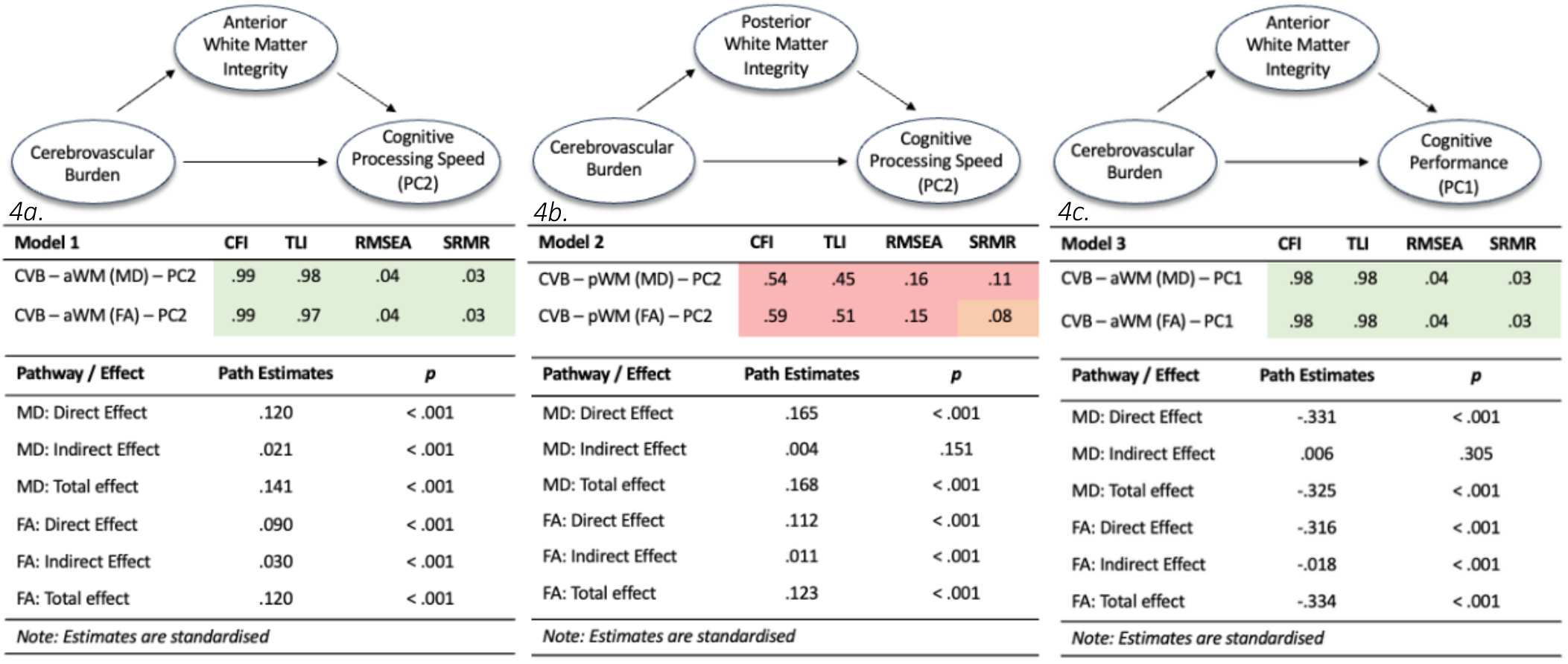
Structural equation models, path loadings and model fit indices. Each figure exhibits a SEM path diagram, standardised path estimates and model fit indices for each of the three models – baseline model, posterior WMI model & general cognitive performance model, respectively. Models were deemed to be a good fit based on the following cut off points: .90 or above (CFI & TLI) or .08 or below ( RMSEA & SRMR). Values highlighted in green indicate good fit, red indicate bad fit and orange indicates borderline. CFI – comparative fit index, TLI – tucker lewis index, RSMEA – root mean square error of approximation, SRMR – standardised root mean square residual.

### Establishing the regional specificity

Regional specificity of the relationship between WMI and the processing speed latent construct was tested by replacing anterior WMI with posterior WMI. In this new model the mediation effect was either no longer significant (MD) or remained significant but reduced (FA), relative to the baseline model (*Figure 4b*). The model containing posterior WMI also showed a significantly poorer fit, when compared with the baseline model (*p* < .001; see *Figure 4b*).

### Establishing the cognitive specificity

To test the cognitive specificity of this effect to processing speed, the outcome measure of the processing speed principal component was replaced with the general cognitive performance principal component. The direct effect between cerebrovascular burden and cognitive outcome remained significant, showing that greater cerebrovascular burden was associated with worse cognitive performance. However, the indirect effect was either no longer significant (MD) or remained significant but reduced (FA), as shown in *Figure 4c*. This demonstrates an inconsistent and weak mediatory effect of anterior WMI between cerebrovascular burden and general cognitive performance. Model fit indices demonstrated good fit, statistically comparable to the fit of the baseline model – which was expected, given that only one variable had changed from the baseline model.

### Multiple Regression Analyses

A multiple regression analyses was conducted to assess whether accounting for risk factor duration can improve the ability for cerebrovascular risk factors to predict cognitive processing speed change. In model 1, *‘smoking’* and *‘diabetes’* reflect binary variables, indicating whether participants are current/past active smokers or if they had previous history of diabetes. In model 2, *‘smoking’* reflects the number of years participants have smoked (pack years) and *‘diabetes’* reflects the number of years since participants received a diabetes diagnosis, where relevant. Model 2 also has an added variable of ‘*hypertension duration*,*’* indicating the number of years since participants received a hypertension diagnosis. Sex, Townsend deprivation index, and ethnicity were also included in both models as covariates. A smaller subset of the sample (N = 22,475) was used in the regression analysis, as risk factor duration data was not available for a proportion of the sample who had indicated they had a given risk factor.

### Overall Model Effects

Model 1, containing static risk factor information, demonstrated significant effects (*F(*29, 22475) = 33.62, *p* < .001, *R*^*2*^ .042), highlighting the ability of cerebrovascular burden to predict processing speed performance. Model 2, which replaced static risk factors for ‘*Smoking’, ‘Diabetes’* with risk factor duration data and added ‘*Hypertension Duration’*, also demonstrated significant effects (*F(*30, 22474) = 34.51, *p* < .001, *R*^*2*^ .044). The difference between the two models was significant, with model 2 explaining a greater amount of variance, relative to model 1 (ANOVA; F(1, 22475) = 57.89 , *p <* .*001)*. This suggests risk factor duration explains a unique proportion of the variance and improves the ability of cerebrovascular burden to predict processing speed performance.

### Individual Risk Factor Effects

All variables, apart from *diabetes* in both model 1 and model 2 were significant predictors of processing speed performance (as derived from *Principal Component 2*), each explaining unique variance. The largest amount of variance was explained by *Cholesterol* in both model 1 (*β*= .033, σ = 0.003) and model 2 (*β* = .028, σ = 0.003). This was followed closely by both *systolic* (model 1 - *β*= .026, σ = 0.002; model 2 - *β* = .021, σ = 0.002) and *diastolic* (model 1 - *β*= -.021, σ = 0.002; model 2 - *β* = - .021, σ = 0.002) blood pressure. The lowest significant contributors in the model were *Smoking* (model 1 - *β*= .007, σ = 0.002; model 2 - *β* = -.003, σ = 0.001) and *BMI* (model 1 - *β*= -.008, σ = 0.001; model 2 - *β* = -.009, σ = 0.001). The regression coefficients for each risk factor in each model are shown in *Figure 6*.

**Figure 6.**
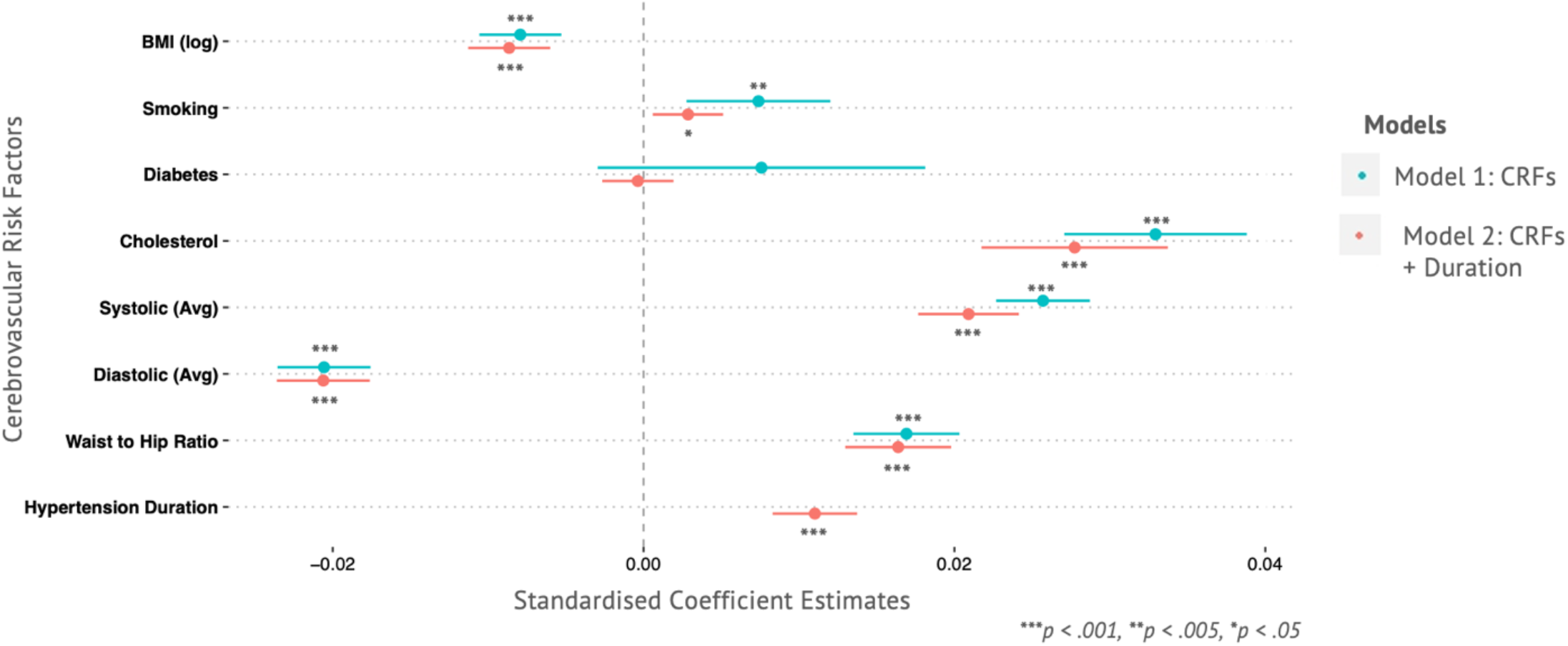
Cerebrovascular risk factors (model 1) and risk factor duration (model 2) predicting processing speed performance. Model 2 depicts the relationship between individual cerebrovascular risk factors and processing speed. The second replicates model 1, with the addition of risk factor duration for Hypertension, Diabetes and Smoking. Note: ‘Diabetes’ in model 1 is a binary yes/no variable based on diagnosis, but in model 2, this reflects how long (in years) participants have had a diagnosis in years. Similarly, ‘Smoking’ in model 1 is binary - indicates whether participants are current or past smokers / never smoked and model 2 reflects number of pack years.

**Figure 7. a (top) & 7b (bottom).**
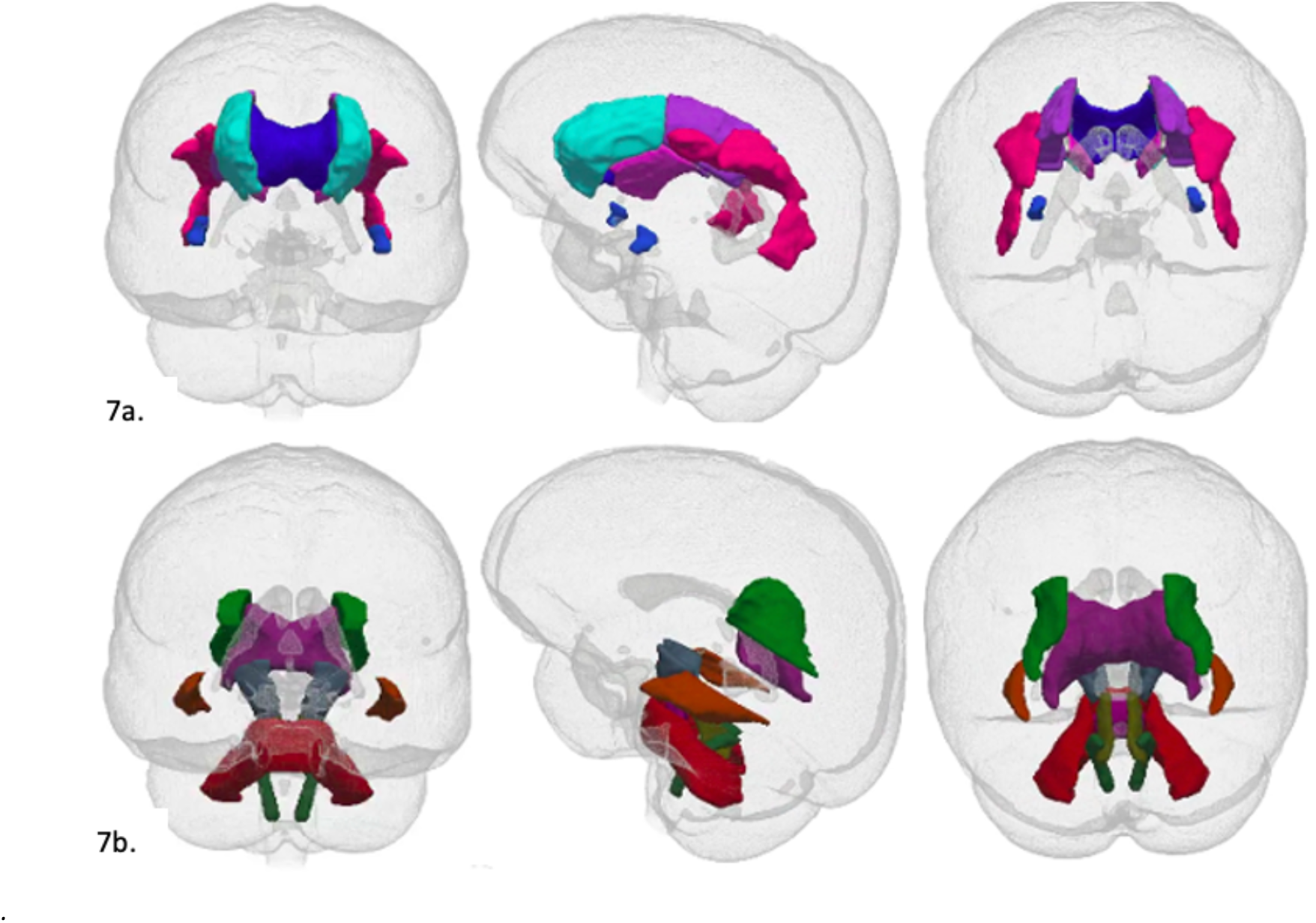
Anterior & posterior white matter tracts. 7a. & 7b. reflect the set of anterior tracts used to create the anterior and posterior white matter integrity latent constructs, respectively.

In model 2 hypertensive duration was found to be a significant predictor, and comparable to established risk factors, such as BMI. In contrast replacing static measures of diabetes or smoking with duration measures had less (non-significant) effect. The findings suggest that accounting for risk factor duration, primarily hypertension duration, significantly improves the prediction of processing speed from cerebrovascular burden, relative to static burden models that do not account for risk factor duration.

## Discussion

In a large sample of people without current or previous neurological disease, we found that cerebrovascular burden was associated with alterations in cognitive function and regionally specific white matter pathology. More specifically, we found that increased cerebrovascular burden was associated with slowed processing speed – and that this effect was mediated by anterior WMI. The mediatory role of WMI was either diminished or no longer significant when anterior tracts were replaced with posterior tracts, or when processing speed was replaced with a metric of general cognitive performance. Together, these findings demonstrate the regionally specific mediatory effect of anterior WMI on the relationship between cerebrovascular burden and processing speed. In addition, we demonstrate the relative impact of cerebrovascular risk factors and risk factor duration on cognitive processing speed change. Each individual cerebrovascular risk factor, except diabetes, significantly predicted slowed processing speed, except for diabetes – which may be reflected by the absence of data to separate type 1 and type 2 diabetes in the UK Biobank. In addition, accounting for risk factor duration for hypertension, smoking and diabetes, improved predictions of processing speed and explained unique variance. These findings support our initial predictions, that a mechanistic pathway, through which cerebrovascular risk factors impact cognitive processing speed, is selectively mediated by alterations in anterior WMI.

Our findings add to the spectrum of vascular diseases and risk factors linked to heightened susceptibility of anterior white matter pathology, such as atrial fibrillation ^28^, subcortical ischaemic vascular disease, ^29^ vascular dementia ^30^ and cerebrovascular risk factors ^15 31^. The pattern of anterior-focused white matter change shown, is also notably similar to the regional ischemic vulnerability of the brain to hypoperfusion shown by clinical stroke samples ^32^. While the mechanisms underpinning this regionally specific vulnerability are incompletely understood, it likely has a vascular basis ^33^. One proposed mechanism is that anterior regions are particularly susceptible to haemodynamic derangement, whereby the vasculature undergoes structural changes and arterioles become tortuous or coiled ^30 34^ – and further exacerbated by cerebrovascular burden ^35^. Consequently, this likely reduces local blood flow and limits white matter perfusion, leading to the emergence of white matter pathology ^36^. However, further work is required to unpick the cellular, haemodynamic, and vascular mechanisms underpinning the relationship between cerebrovascular burden and anterior white matter alterations.

We also demonstrate the cognitive consequences - particularly for processing speed - of cerebrovascular burden via breakdown of anterior WMI. This link between anterior WMI and slowed processing speed is consistent with findings from a range of white matter-affected diseases, such as multiple sclerosis ^37^, Alexander disease ^38^ and cerebral small vessel disease ^20^. There are several mechanisms that likely drive this affect, such as myelin loss, axonal integrity alterations and other white-matter pathological processes indicated by various diffusion metrics. These factors likely prevent, or slow, neural signalling, ultimately resulting in slowed cognitive processing speed ^39^. This view is further supported by previous associations between the presence of highly myelinated tracts and faster processing speed ^40^. We also demonstrated a link between cerebrovascular burden and more general cognitive performance, which was not well explained by anterior WMI. Different white matter networks may be impacted by cerebrovascular burden and may better explain changes in other cognitive domains and highlights the preferential contribution of anterior WMI to processing speed performance.

Our findings offer practical suggestions to improve cerebrovascular burden quantification. Traditional methods of cerebrovascular risk quantification used in clinic and research assess relative cerebrovascular burden from a static, cross-sectional viewpoint and commonly use methods such as the Framingham Risk score ^41^ and QRISK ^42^. Here we show, for the first time in one model, that modelling risk factor duration, specifically for hypertension and diabetes duration and smoking pack years, can significantly improve the extent to which cerebrovascular burden predicts processing speed. These findings suggest traditional methods may underestimate the extent of cerebrovascular burden in health and disease. Although risk factor duration metrics were participant-estimated, our findings align well with previous work. For example, smoking duration and intensity has been found to negatively predict cognitive function ^43^. In addition, previous work used clinic-confirmed hypertensive duration metrics to highlight a negative relationship with cognitive function ^25^. These findings converge to suggest that participant estimates of risk factor duration usefully enhance predictions of cerebrovascular burden on cognitive function. Therefore, including this information, where available, will lead to more sensitive predictive models of brain health and cognitive function.

### Limitations & Future Research Directions

Despite the large-scale sample and quality of the UK Biobank data, there are several methodological limitations to consider. First, we used cross-sectional data, which is limited by several caveats – including the inability to confidently claim causal inference. However, the inclusion of risk factor duration metrics in our work provides an estimate of longitudinal insight into the body-brain-cognition relationships described, indicating burden builds over time for individual risk factors. The schedule of re-imaging currently underway by the UK Biobank will allow longitudinal analyses and complement our findings on cerebrovascular change and cognition. Second, our work cannot be used to determine the optimal time windows for risk factor alleviation. While we demonstrate the impact of cerebrovascular burden on both brain integrity and cognition, many risk factors used in these analyses are modifiable. Previous work has suggested a limited window in which alleviation of cerebrovascular risk factors can yield a beneficial impact on both brain integrity and cognitive function, especially in later life ^44 45^. This window is likely defined by the interaction between the degree of burden and age, but to date this relationship is poorly characterised in the literature, despite its potentially critical role in reducing the likelihood of dementia. Third, those who opted to participate in the UK Biobank tend to represent less socioeconomically deprived areas, with fewer morbidities than the rest of the UK, with comparatively lower rates of obesity and smoking ^46^. Although this may at least slightly limit the generalisability of the results, our findings were derived from a relatively healthy population, suggesting that these effects may in fact be exacerbated in a more representative sample.

In summary, our work proposes a mechanistic pathway through which cerebrovascular burden impacts brain integrity and cognitive function, providing evidence of both regional and cognitive specificity. We show that greater cerebrovascular burden was associated with slowed processing speed – an effect at least partially mediated by alterations in anterior WMI. We demonstrate that models accounting for risk factor duration can improve predictive outcomes. These findings are shown in a large, generally healthy cohort, where vascular insult is relatively mild, and the sample is primarily middle aged. Addressing these risk factors as early as possible will likely aid in the prevention of later-life clinically significant cognitive decline. Through future work we can establish the pace at which these structural and cognitive changes occur – and the time windows during which effective interventions can optimally be applied.

## Methods

### Participants

Participant’s data were obtained from the UK Biobank’s latest release, as of March 2021. Participants with missing cognitive, cerebrovascular and neuroimaging data were excluded. In addition, participants were excluded if they presented with a history or current diagnosis of neurological disease, head/brain injury or trauma, stroke, TIA, alcohol or opioid dependence, Epilepsy, Parkinson’s disease, Alzheimer’s disease, Dementia, cognitive impairment, chronic or degenerative neurological issues (including demyelination-associated syndromes), brain hematoma, brain abscess, meningitis, encephalitis, infection of the nervous system, haemorrhage or multiple sclerosis during their medical interview. As outlined in the non-cancer illness codes (https://biobank.ndph.ox.ac.uk/showcase/coding.cgi?id=6). In total, data from 37,265 participants were used in the main analysis, aged 40-70. A smaller subset of the sample (N = 22475) was used in the regression analysis, due to missing risk factor duration data.

### Cognitive Testing

Computerised cognitive testing was administered using a touch-screen during the MRI visit. The cognitive tests analysed were tests of numeric memory, fluid intelligence, trail making (numeric and alphanumeric), matrix pattern completion, symbol digit substitution, tower rearranging, paired associative learning, pairs matching and a reaction-time based game of snap. Detailed summaries of each individual cognitive assessment undertaken during the neuroimaging visit can be found elsewhere and in UK Biobank documentation (https://biobank.ctsu.ox.ac.uk/crystal/label.cgi?id=100026).

### MRI Data Acquisition & Analysis

Imaging data for the Biobank were collected using a Siemens Skyra 3T Scanner and 32-channel head coil, per the openly available protocol and acquisition detail documentation (https://biobank.ctsu.ox.ac.uk/crystal/crystal/docs/brain_mri.pdf & https://www.fmrib.ox.ac.uk/ukbiobank/protocol). In short, the dMRI acquisition consisted of a 2mm isotropic spin-echo multiband echo-planar sequence, with 50 *b* = 1000s mm and 50 *b* = 2000s mm diffusion weighted volumes (100 encoding directions total).

All imaging data were quality checked internally by the UK Biobank team, before being processed by an openly available automated pipeline^47^. One component of this pipeline is the generation of image-derived phenotypes (IDPs), which are individual tract or volumetric averages of specific regions in the brain. The IDPs of interest, used in the current analysis were tract-averaged fractional anisotropy (FA) and mean diffusivity (MD) metrics for the following anterior white matter tracts: genu of corpus callosum, anterior corona radiata, superior corona radiata, anterior limb of internal capsule, superior fronto-occipital fasciculus, superior longitudinal fasciculus, uncinate fasciculus (as shown in *Figure 7a)*. The control, posterior tracts used were cerebral peduncle, inferior cerebellar peduncle, middle cerebellar peduncle, superior cerebellar peduncle, posterior corona radiata, splenium of corpus callosum, sagittal stratum and pontine crossing tract (shown in *Figure 7b)*.

**Figure 8.**
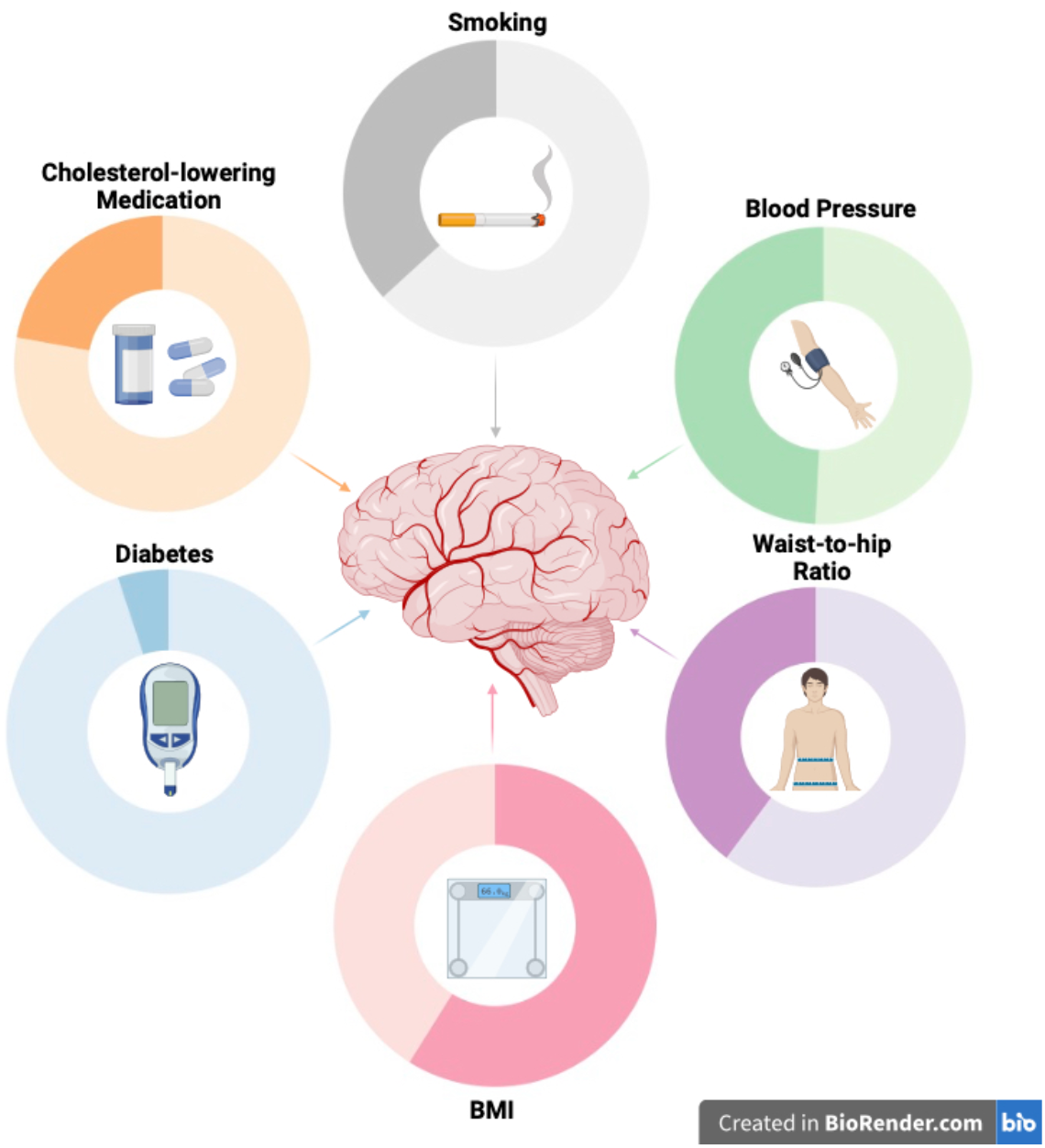
Prevalence of individual cerebrovascular risk factors in the sample. Figure 8 depicts the proportion of the sample with each individual risk factor – indicated by the darker proportion of each circle. To be considered to have a given risk factor, they needed to meet the following criteria for each risk factor – smoking: indication of current or past active smoking status, blood pressure: 140/90mmHg, waist-to-hip ratio: females - < .80, males - <.90, BMI: >25, Diabetes: diagnosis confirmed by a doctor, cholesterol: currently taking cholesterol lowering medication. Figure was created used BioRender.

### Cerebrovascular Burden

Cerebrovascular risk factor information was collected from participants during a medical history interview, during their neuroimaging session. Where continuous variables were unavailable, binary risk factors were coded based on the presence of a given risk factor, described in detail below. *Figure 8* depicts the proportion of the sample with any given risk factor.

#### Smoking

Participants who indicated current or past active smoking status were coded with a score of 1, those who did not were coded with a score of 0.

#### Diabetes

Participants who indicated that they had received a diabetes diagnosis were coded with a value of 1 and those who had not, were coded with a value of 0.

#### Hypercholesterolaemia

A current prescription of cholesterol-lowering medication was used as a surrogate measure for high cholesterol and participants who indicated that they take cholesterol-lowering medication were coded with a value of 1 and those who did not, a value of 0.

#### Blood Pressure

Blood pressure (BP) was measured twice, moments apart using an automated Omron digital BP monitor for both diastolic and systolic BP, which were then averaged for each.

#### Waist-to-hip ratio & BMI

Anthropometric measures were taken manually once participants had removed any bulky clothing. Waist and hip circumference measures were transformed into a ratio metric by dividing the waist hip measurements. BMI was calculated by dividing weight (kg) by the square of height in metres.

#### Risk Factor Duration

Risk factor duration was calculated for each risk factor, where available, specifically - hypertension, smoking and diabetes.

#### Hypertensive duration

Participants who had indicated that they were currently on hypertensive medication were asked to estimate the age they were diagnosed with high blood pressure, which was subtracted from their current age.

#### Smoking duration

To quantify how long each participant with a positive smoking status had been smoking, pack years was calculated by dividing the number of cigarettes smoked per day by 20, multiplied by the number of years smoking.

#### Diabetes duration

Participants who indicated that they were diabetic were asked to estimate the age they were diagnosed with diabetes, which was subtracted from their current age.

### Statistical Analyses

Firstly, data were visually inspected to assess the distribution of each variable and to ensure they met the necessary parametric assumptions. Participants with IDPs = 0 were omitted entirely from the dataset. BMI data were log-transformed, to correct for skewed distribution. Any participants with missing cerebrovascular risk, cognitive or neuroimaging were omitted from the dataset & analysis.

A principal component analysis (PCA) was performed to derive two new latent constructs from all cognitive data available from the UK Biobank during the participants’ imaging visit. With the exception of data from pilot tests, tests with significant amounts of missing data and those unavailable for download were not included. Due to the volume of missing cognitive data from the UK Biobank, the ‘*MissMDA’* package was used to obtain scores and loadings ^48^. This package allows PCA to be performed on datasets with missing values, by using an iterative PCA algorithm to impute missing data and perform the PCA. We then used the factor loadings from PC1 and PC2 in our SEMs. Based on the factor loadings, we described PC1 to reflect more general cognitive performance, as tasks such as associative memory, tower rearranging, matrix pattern completion and fluid intelligence loaded highly, which all measured performance. We described PC2 to represent cognitive processing speed, as the cognitive tests trail making, pairs matching and snap loaded highly, which were measures of speed.

In preparation for the SEM, a latent construct of cerebrovascular burden was created, loaded with each cerebrovascular risk factor using factor analysis. This method of burden quantification has been validated in previous studies and provides a more informative indicator of body-brain relationships, relative to commonly used aggregate quantification methods ^14 15^. Similarly, two latent constructs of anterior WMI were also created, with left & right tracts loaded separately for each construct - one representing FA and the other, MD.

In the SEM, the latent construct of cerebrovascular burden (loaded as cerebrovascular risk factors, as described) acted as the predictor variable. The latent construct of anterior WMI acted as the mediator variable. Two separate models were created - one containing FA and the other, MD. The principal component for processing speed was used as the outcome variable for the baseline SEM. The ‘*lavaan 0*.*6-12’* package ^49^ was used to build and analyse the SEMs.

To establish the regional specificity of this effect, a latent construct of posterior WMI was also created (see *Figure 1b)*. A new SEM was built, identical to the baseline model, but the anterior WMI mediatory variable was replaced with the posterior WMI latent variable. To explore cognitive specificity, a third SEM was built, but here the outcome variable was replaced with the principal component for general cognitive performance, rather than processing speed.

A multiple regression analysis was also performed to characterise the relative relationship between individual risk factors, risk factor duration and processing speed. All cerebrovascular risk factors were used as predictor variables and processing speed acted as the outcome variable, with ethnicity, Townsend Deprivation Index & sex added in as covariates to the model. To assess whether accounting for risk factor duration can improve the predictive utility of cerebrovascular burden on processing speed, the smoking status variable was replaced by smoking duration, diabetes duration replaced diabetic risk and hypertensive duration was added into the multiple regression model.

## Acknowledgements

We thank members of the Memory Research Unit, GJBRC & MISC Lab for their feedback and support throughout this project.

## Author contributions

KLM was involved in study conception, design, data analysis and interpretation and writing of the manuscript. EM contributed to the data analysis and commented on the manuscript. SA and CS contributed to data interpretation and commented on the manuscript. DM and NM conceptualised, designed and managed the study, and contributed to the data interpretation and writing of the manuscript.

## Data availability statement

All UK Biobank information is available online on the webpage www.ukbiobank. Data access is available through applications.

